# Engineering of a Polydisperse Small Heat-Shock Protein Reveals Conserved Motifs of Oligomer Plasticity

**DOI:** 10.1101/247288

**Authors:** Sanjay Mishra, Shane A. Chandler, Dewight Williams, Derek P. Claxton, Hanane A. Koteiche, Phoebe L. Stewart, Justin L.P. Benesch, Hassane S. Mchaourab

## Abstract

Small heat-shock proteins (sHSP) are molecular chaperones that bind and sequester partially and globally unfolded states of their client proteins. Of paramount importance to their physiological roles is the assembly into large oligomers, which for mammalian sHSP are polydisperse and undergo subunit exchange. The flexibility and dynamic nature of these oligomers mediates functional regulation by phosphorylation and underpins the deleterious effects of disease-linked mutations. Previously, we discovered that the archaeal Hsp16.5, which natively forms ordered and symmetric 24-subunit oligomers, can be engineered to transition to an ordered and symmetric 48-subunit oligomer by insertion of a peptide from human HspB1 (Hsp27) at the junction of the N-terminal and α-crystallin domains. Here, we carried out a detailed analysis of the determinants of Hsp16.5 oligomeric plasticity by altering the sequence and length of the inserted peptide. Utilizing light scattering, blue native gel electrophoresis, native mass spectrometry and electron microscopy, we uncovered the existence of an array of oligomeric states (30 to 38 subunits) that can be populated as a consequence of different insertions. These oligomers are intermediate states on the assembly pathway of the 48-subunit oligomer as two of the variants can concurrently form 24-subunit or 30-38 subunit polydisperse oligomers. Polydisperse Hsp16.5 oligomers displayed higher affinity to a model client protein consistent with a general mechanism for recognition and binding that involves increased access of the hydrophobic N-terminal region. Our findings, which integrate structural and functional analyses from evolutionarily-distant sHSP, support a model wherein the modular architecture of these proteins encodes motifs of oligomer polydispersity, dissociation and expansion to achieve functional diversity and regulation.

## Introduction

Small heat-shock proteins (sHSP) are a ubiquitous class of molecular chaperones that play a central role in the stress resistance of all organisms^1,2^. While they share the general property of binding non-native and unfolded proteins with other heat shock proteins, sHSP chaperone activity does not involve the direct consumption of ATP^3,4^. They bind their client proteins in stable complexes thereby inhibiting aggregation and as such provide an energy-efficient, first line defense against protein aggregation^1,5,6^. Humans express 10 distinct sHSP that have critical physiological roles including maintenance of lens transparency and the integrity of cardiac and skeletal muscles^7–11^. Mutations in human sHSP have been associated with inherited diseases, most notably cataract and cardiomyopathy^12–17^.

The molecular architecture of sHSP is built around a highly conserved 90-amino acid domain, the α-crystallin domain^11,18^. This domain forms a dimeric building block which assembles into oligomers through interactions in the highly variable N-terminal domain and a C-terminal tail^19–28^. Crystal structures and spectroscopic analyses have defined the architecture of this building block across the evolutionary spectrum, identifying two motifs of dimerization so far^21,23,24,29–31^. The packing of these dimers dictates the structure of the sHSP oligomers which display remarkable diversity in size and symmetry^32^.

Far from being static “sponges” for unfolded proteins, sHSP oligomeric structures, particularly from eukaryotes, are characterized by a fascinating spectrum of conformational flexibility^1,33,34^. Multiple studies have demonstrated that polydispersity of sHSP oligomers is a mechanism for regulation of binding affinity and capacity^35–39^. Dynamic oligomer dissociation, manifested by subunit exchange between oligomers, exposes a flexible N-terminal region containing putative interaction sites that are otherwise inaccessible^40,41^. Shifts in the equilibrium between large oligomers and dimers, e.g. by phosphorylation of Hsp27 and αB-crystallin, enhance the affinity to client proteins and the binding capacity^42–44^. Alternatively, oligomer expansion resulting from an increase in the number of subunits per oligomer can expose the N-terminal region^45,46^. Initially observed in cataract-linked mutants of α-crystallins, the structural basis of expansion was uncovered by an engineered variant of *M. Jannaschii* Hsp16.5^47,48^. Insertion of a 14-amino acid peptide (hereafter referred to as the P1 peptide), found at the junction of the N-terminal and α-crystallin domain of human Hsp27 (HspB1), at the equivalent position in Hsp16.5 resulted in the expansion of the oligomer from 24 subunits to 48 subunits^45,47,48^. A high resolution structure of the expanded Hsp16.5 variant revealed that the changes in the sequence of the N-terminal region lead to two different conformations of the C-terminal tail which in turn enable two different packing interactions of the α-crystallin dimer within the same oligomer^45^.

The expansion of the P1-containing Hsp16.5 variant was unexpected considering the lack of polydispersity of WT Hsp16.5, and raised the question of whether oligomeric plasticity is in an intrinsic property of sHSP. That is, are all sHSPs capable of displaying a broad spectrum of oligomeric structures? A related question is whether the P1 peptide sequence encodes specific structural and dynamic properties that are essential to the observed oligomer expansion. While P1 is not native to Hsp16.5, its role in modulating oligomer dissociation of human Hsp27 implies evolutionary optimization of its sequence motifs^36,41,48^.

The work reported here follows up on the structural and functional studies of the expanded Hsp16.5, hereafter referred to as Hsp16.5-P1, by investigating the sequence determinants in the P1 peptide that drive expansion of the oligomer. For this purpose, the amino acid composition and/or length of the P1 peptide sequence were manipulated (Figure 1) and the resulting variants of Hsp16.5 were characterized for the size of the oligomer and its polydispersity by size exclusion chromatography (SEC), multi-angle light scattering (MALS), and blue native gel electrophoresis (BN-PAGE). Furthermore, we visualized the various possible oligomeric assemblies using negative stain electron microscopy (EM) and analyzed oligomer distributions by native mass spectrometry (MS)^49^. Finally, the changes in the oligomeric assembly of the variants were correlated with their affinities to a destabilized mutant of T4 lysozyme.

**Figure 1.**
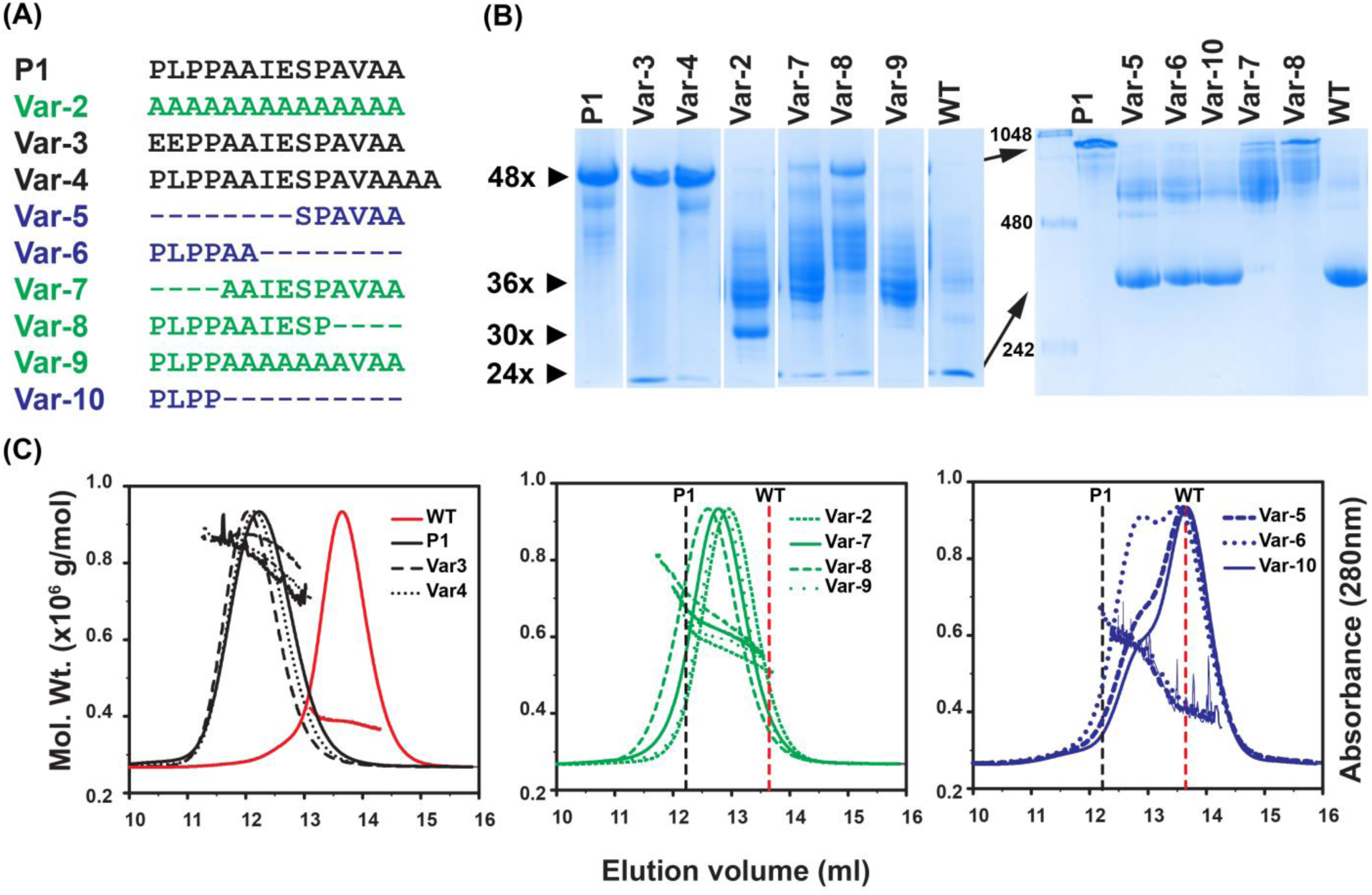
Characterization of the oligomers formed by insertion of peptides in Hsp16.5. A, Sequences of the peptides inserted in the Hsp16.5 sequence between residues 33-34. The variants are grouped by color according to oligomeric properties shown in panels B and C. B, BN-PAGE. Left panel, 5% acrylamide gel; right panel, 7.5% acrylamide gel. The approximate number of subunits corresponding to band mobility is shown with small arrows on the left side of the left panel. The molecular weight marker is shown in the first lane of the right panel. Large arrows highlight mobility differences of the P1-insert and WT constructs in the 7.5% acrylamide gel. C, SEC-MALS analysis of purified variants. Variants are grouped according to similar characteristics. The molecular mass range that is sampled across the elution peak is indicated by the slope of the line for each variant. The vertical lines benchmark the WT (dotted red) and the P1-insert (dotted black) elution peak.

A number of general themes emerge from this systematic investigation that extends the current understanding of the structural basis of sHSP polydispersity. We find that minor changes in the amino acid sequence of P1 lead to previously unobserved oligomeric assemblies characterized by numbers of subunits distinct from the WT and P1 forms. Two of the engineered variants can form 24- and 36-subunit oligomers, suggesting that the latter are intermediate states on the assembly pathway from 24 to 48 subunits. Moreover, a sequence motif in P1 is identified as critical to the dynamic oligomer dissociation and thus the chaperone activity of human Hsp27. Finally, analysis of substrate binding affinities of the variants further reinforces the relationship between oligomer size, polydispersity and chaperone efficiency.

## Results

### Changes in the sequence of P1 enhance the polydispersity of Hsp16.5-P1

Inspection of the P1-peptide reveals three distinct sequence characteristics: a PLPP sequence at the N-terminus and an alanine rich AVAA sequence at the C-terminus interrupted by an IESP sequence (Fig. 1A). To define the contribution of each of these elements to the expansion of the oligomer, we generated nine new variants (Fig. 1A), that were expressed in *E. Coli* (Fig. S1), and then analyzed for oligomer size and distribution by BN-PAGE (Fig.1B) and SEC in conjunction with MALS (Fig. 1C). We found, in accordance with previous results, that insertion of the native P1 peptide leads to a well-defined expanded 48-subunit oligomer with an average molar mass of ~0.8 MDa as observed by SEC-MALS (Fig. 1B-C and Table 1)^45,47,48^. Mutation of two N-terminal residues of P1 to acidic residues (Var-3) or extension of P1 length by two Ala residues at the C-terminus (Var-4) did not change the average 48-subunit oligomeric assembly. However, the molar mass distribution of Var-4 revealed a more polydisperse ensemble of oligomers and BN-PAGE analysis indicated the presence of a 24-subunit oligomer more prominently for Var-3. Increasing the redundancy of the peptide sequence by complete substitution with an isometric poly-Ala insert (Var-2) leads to oligomers smaller than the Hsp16.5-P1 but larger than Hsp16.5 (Fig. 1C). The average molar mass of Var-2 corresponds roughly to an assembly of 36 subunits and represents a previously unobserved oligomer size (Table 1).

**Table 1.** Average molar mass and average hydrodynamic radius moments of the HSP variants

| HSP | Average<br>Molar mass<br>($\times 10^6$ g/mol) | Average<br>Hydrodynamic<br>radius (nm) |
| --- | --- | --- |
| P1 | 0.80 | 8.2 |
| Var-2 | 0.58 | 7.0 |
| Var-3 | 0.86 | 8.3 |
| Var-4 | 0.84 | 8.2 |
| Var-5<sup>♦</sup> | 0.53 | 7.0 |
|  | 0.40 | 6.2 |
| Var-6<sup>♦</sup> | 0.59 | 7.2 |
|  | 0.44 | 6.4 |
| Var-7 | 0.64 | 7.4 |
| Var-8 | 0.69 | 7.7 |
| Var-9 | 0.60 | 7.4 |
| Var-10<sup>♦</sup> | 0.56 | 7.1 |
|  | 0.40 | 6.1 |
| WT | 0.39 | 6.1 |

The molar masses and the radii were measured by the multi-angle light scattering of the proteins eluted from the size exclusion chromatography, using Astra software (Wyatt). ♦These variants segregate into two distinct populations.

### Shortening the P1 peptide stabilizes a 36-subunit oligomeric cluster

The ensemble of oligomers clustered around the 36-subunit species also appeared in other variants of shorter length. Truncation of the P1-peptide, either from its N-terminus (Var-7), or the C-terminus (Var-8) by 4 residues resulted in oligomer assemblies of average size approximating 36 subunits similar to Var-2. Despite a comparable average molar mass, the oligomeric populations of Var-7 and Var-8 were more polydisperse than Var-2, as reflected by larger spreads in molar mass distributions across their SEC profiles (Fig. 1C). Remarkably, further truncation of the P1-peptide by an additional four residues from either the N-terminus (Var-5) or C-terminus (Var-6) leads to two distinct populations corresponding to a 36-subunit cluster and WT-like 24-subunit oligomers (Fig. 1C). The concurrent assembly into two forms suggests that they are of similar free energy. The relative population of the 36-subunit oligomer was greater in the Var-6 construct where the N-terminal half of P1 remained intact. Similarly, a shift toward formation of larger oligomeric species was observed in Var-8 where the N-terminus of P1 remains intact relative to Var-7 where the first 4 amino acids where truncated (Table 1).

Comparative analysis by BN-PAGE (Fig. 1B) was consonant with the relative polydispersity within the oligomeric ensemble of the variants observed by SEC-MALS. WT Hsp16.5 and Hsp16.5-P1 resolved as near homogenous 24- and 48-subunit entities, respectively, on a 7.5% acrylamide gel. BN-PAGE (5% acrylamide gel) of Var-3 and Var-4 also confirmed assembly predominately into 48-subunit oligomers. In contrast to these defined oligomers, Var-2, Var-7 and Var-8 demonstrated broad heterogeneity around the putative 36-subunit oligomer in agreement with the molar mass distributions inferred from SEC-MALS. Importantly, BN-PAGE exposes a discrete oligomer species for Var-2 which appears to consist of about 30 subunits.

These observations, demonstrating that P1’s chemical properties and length modulate the oligomer size distribution of Hsp16.5, suggest that this sHSP is intrinsically amenable to polydispersity. Furthermore, the N-terminus of the P1 sequence appears to play a more prominent role in promoting formation of larger oligomers. Indeed, insertion of only the four leading N-terminal PLPP residues (Var-10) was sufficient to generate a 36-subunit cluster. We suggest that the 36-subunit cluster represents assembly intermediates along the Hsp16.5 expansion pathway.

### EM analysis of the 36-subunit cluster

While SEC-MALS and BN-PAGE revealed changes in oligomer size and distribution, they are intrinsically low resolution indicators of heterogeneity. Therefore, to visualize the structural features of the previously unobserved oligomeric assemblies, we analyzed the variants by negative stain EM as shown in Fig. 2 and Fig. S2. The particles were classified broadly into three classes: a WT-like 24-subunit oligomer, a 36-subunit oligomeric cluster, and the 48-subunit expanded form (Table 2). A fourth minor class is that of a 30-subunit oligomeric cluster. The observed overall trends were consistent with the SEC-MALS and BN-PAGE analysis. Thus, the insertion of the P1 peptide into Hsp16.5 expanded more than 90% of particles into the 48-subunit oligomer. Changes in P1 sequence and length induced substantial shifts in the populations of each species. Unlike the native P1 sequence, the poly-Ala Var-2 insertions predominantly caused expansion to the 36-subunit cluster, emphasizing that insertion length alone does not determine oligomer size. Furthermore, the Var-3 construct only slightly reduced the population of 48-subunit oligomers, but the addition of two Ala residues to the P1 sequence (Var-4) strongly perturbed the fully expanded oligomer. Notably, sequences shorter than the native P1-peptide were ineffective in maintaining complete expansion of the oligomer. Almost all particles of Var-7 and Var-8 assembled into the 36-subunit oligomer; whereas Var-5 and Var-6 assembled into both the 36-subunit and the 24-subunit oligomers.

**Table 2.** Particle distribution of HSP variants determined by negative stain EM

| HSP variant | 24 subunit | 36 subunit | 48 subunit |
| --- | --- | --- | --- |
| P1 | - | 7.5 | 92.5 |
| Var-2 | 8.5 | 91.5 | - |
| Var-3 | - | 15.1 | 84.7 |
| Var-4 | 3.1 | 57.9 | 39.0 |
| Var-5 | 53.8 | 41.7 | 6.6 |
| Var-6 | 32.9 | 53.1 | 7.4 |
| Var-7 | 12.0 | 86.2 | 3.8 |
| Var-8 | 1.0 | 97.4 | 1.6 |
| Var-9 | - | 100 | - |

The Hsp16.5 mutant datasets were refined against three representative structures of 24, 36, and 48 subunits with the EMAN multirefine subroutine in order to partition individual particle images into one of three classes. For each Hsp16.5 mutant the percentage of particles placed into each of the three classes is shown.

**Figure 2.**
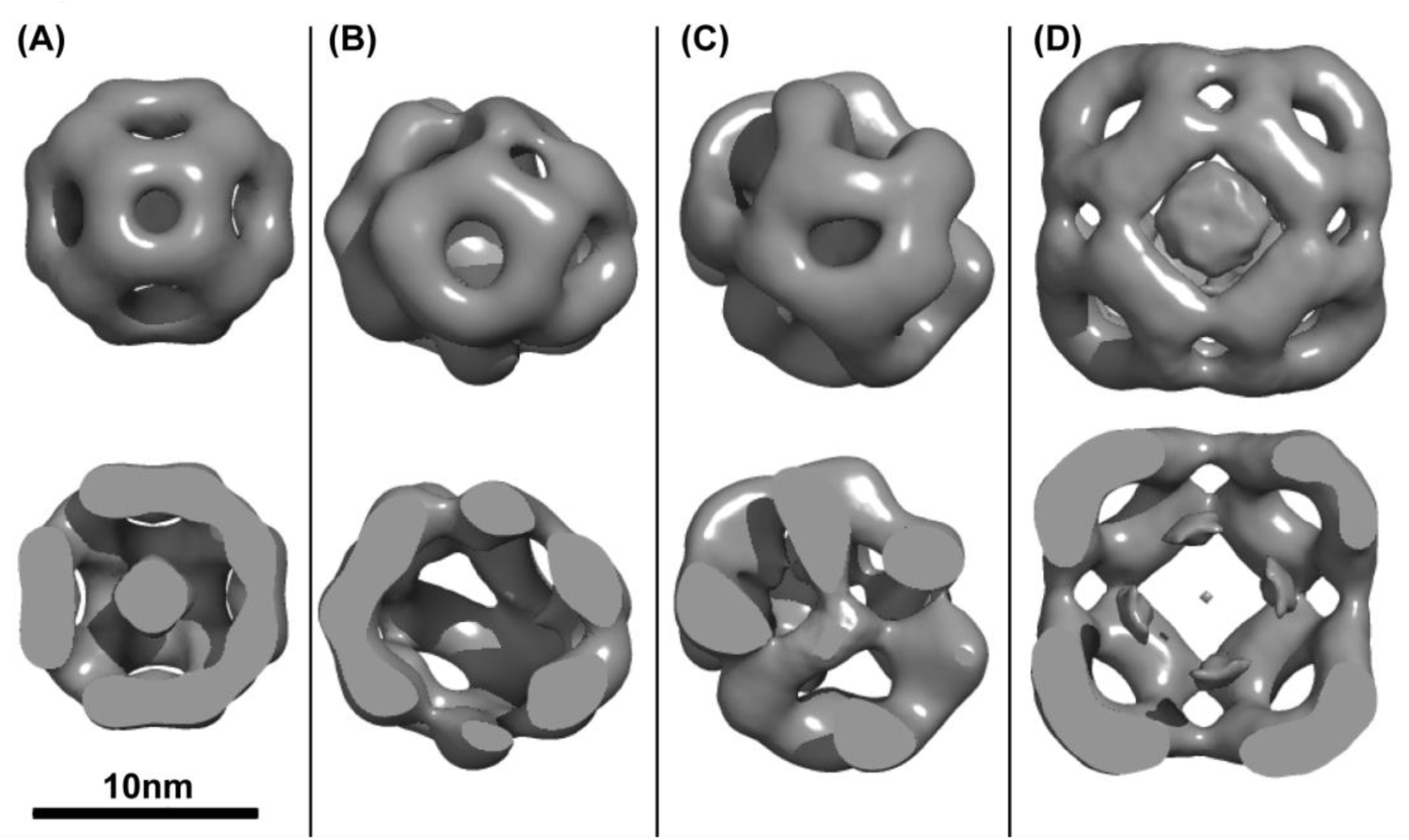
Three dimensional negative stain-EM reconstructions of representative oligomers from peptide insertion variants of Hsp16.5. A, 24-subunit (WT-like). B, 30-subunit. C, 36-subunit. D, 48-subunit (P1-like). Top panel: surface representation; Bottom panel: clipped view. The internal anatomy of the intermediate 30 subunit class structure lacks density whereas the 36 average structure has internal density. This makes the circumference about the same for these two types of particles. In contrast to the well-ordered arrangement of the 24- and 48-subunit oligomers, the intermediate size oligomers demonstrate enhanced heterogeneity and lack of symmetry. The relative particle distribution for each variant is given in Table 2.

The Var-9 insertion, consisting of the N- and C- terminal quartet of P1 sequences with an interceding AAAA sequence, promoted possibly the most homogeneous intermediate assembly centered around 36 subunits (Table 1 and Fig. 1B). Classified Var-9 particles reveal a complex size consistent with a 36 subunit but with both WT and P1 facet arrangement of dimers. The 3D reconstruction from 9824 negative stained particles achieved a modest resolution of 28Å (Fig 2C), which is indicative of a variable assembly. While Var-9 is heterogeneous, it was classified as consisting primarily of 36-subunit oligomers.

3D reconstruction of the Var-6 insertion mutant yielded a 30- subunit assembly. Even after class-based sorting, the resolution of this oligomer was limited to >25Å, suggesting that this intermediate-sized assembly is likewise structurally heterogeneous. The average structures of Var-6 and Var-9 reveal gaps both in the internal assembly of dimers as well as the outer shell that has regions of octahedral-like symmetry. Both the 30-subunit and 36-subunit reconstructions resemble that of the 24-subunit and 48-subunit assemblies, but without complete octahedral symmetry and with regions of incomplete facets. It is likely that even among assemblies with exactly 30 or 36 subunits, there is structural heterogeneity.

Thus, while Hsp6.5 can expand into intermediate size oligomers, the particle images are suggestive of a “molten-globule” like assemblies with incomplete outer shells. Crystallographic analysis of Hsp16.5-P1 demonstrated that introduction of P1 facilitates structural changes in the C-terminal tail which alters its orientation to accommodate oligomer expansion while leaving the α-crystallin dimer unaffected^45^. In the context of this model, the EM analysis of Vars-6 and 9 suggests that either the two configurations of the C-terminal tail corresponding to the WT and P1 oligomer co-exist in the intermediate size assemblies or that this tail adopts a new configuration that is incompatible with either of the two ordered and symmetric oligomers.

### High resolution oligomer binning by native state mass spectrometry

In light of the structural heterogeneity of the 36 oligomer cluster and the consequent low resolution of the EM reconstruction, we analyzed a selected set of variants by native state mass spectrometry to determine the exact stoichiometries populated by the Hsp16.5 variants. All variants examined returned high quality mass spectra featuring signal >8000 m/z, indicative of high-mass oligomers spanning multiple stoichiometries; such dispersity in structure is shown for Var-2 in Figure 3. To compare with EM particle-binning, we extracted the ion counts for the different oligomers, grouped according to 24-, 48-subunits, and stoichiometries in between, by integrating the signal recorded in defined regions of the spectra (Fig S3). The abundances we obtained in this way show a remarkable correspondence with EM data (Table 3), despite the simplifications in this analysis, validating our sorting during single-particle analysis.

**Figure 3.**
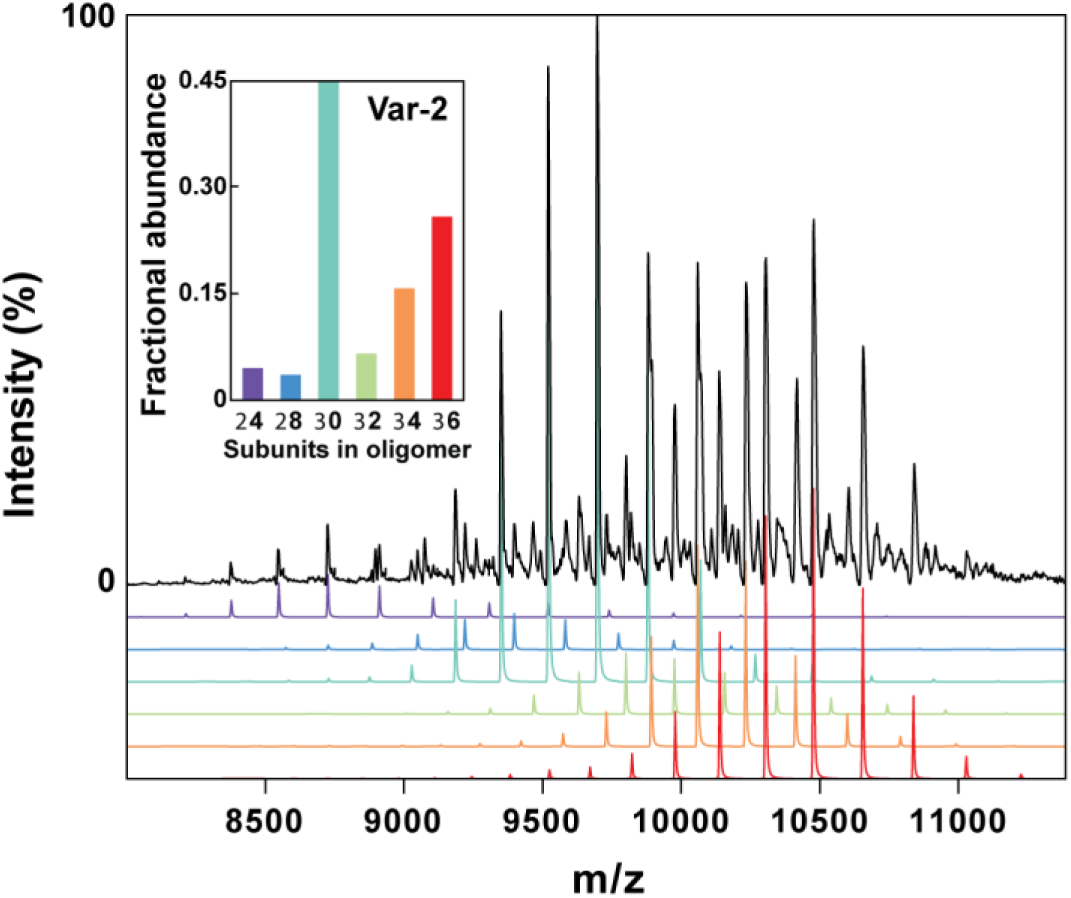
Oligomer binning of Var-2 by native state mass spectrometry. Native mass spectrum of Var-2 revealed the co-population of multiple oligomeric states. The relative abundances of the different states (inset) were extracted from the signal intensity.

**Table 3.** Relative abundances of HSP variants determined by native MS

| HSP variant | 24 subunit | 28-38 subunits | 48 subunit |
| --- | --- | --- | --- |
| P1 | 0.04 | 0.07 | 0.89 |
| Var-2 | 0.05 | 0.91 | 0.04 |
| Var-4 | 0.11 | 0.52 | 0.37 |
| Var-5 | 0.44 | 0.39 | 0.17 |
| Var-6 | 0.69 | 0.24 | 0.07 |
| Var-8 | 0.11 | 0.81 | 0.08 |

The high resolution afforded by the MS data allowed us to identify individual stoichiometries. For all investigated variants, we observed assemblies composed of 28 to 38 subunits, which were separated by dimeric intervals. Using a combination of MS and tandem-MS data, in which the ions were collisionally activated to cause their dissociation and corresponding charge-reduction, we were able to obtain abundances for each individual stoichiometry offering an additional level of detail with good agreement to all methods presented (Figs S4, S5 and Table S1)^49^. This data demonstrated that the 24- and 48- subunit assemblies are well defined entities, with no 26 or 40-46-subunit assemblies observed. In contrast, the 36-subunit assembly was flanked by numerous other assemblies of varying subunit stoichiometries. This observation suggests that the 24- and 48-subunit oligomers are much more favourable symmetries while the 36-subunit is of marginal stability relative to neighbouring stoichiometries. The size heterogeneity of the oligomers is reminiscent of mammalian sHSP. The difference between oligomers by two subunits is consistent with the monomer of Hsp16.5 being highly unstable.

### Stability of the 36-subunit oligomeric state

The observations described above indicate that the α-crystallin dimer-dimer interactions of Hsp16.5 are intrinsically malleable presumably as a consequence of the flexibility of the C-terminal tail. We reasoned that the peptide insertions might also perturb the dynamic properties of Hsp16.5 assemblies, such as subunit exchange between oligomers. Previous studies have shown that Hsp16.5 is thermally stable up to 85°C and maintains a native 24-subunit quaternary structure, even in the presence of rapid subunit exchange at elevated temperatures^29,50–52^. Consistent with these results, exposure to high temperature did not alter the oligomeric state of WT Hsp16.5 or the Hsp16.5-P1 as evident from SEC-MALS profiles (Fig. 4A). Similarly, the Var-7 and Var-8 oligomers, which mostly occupy the 36-subunit intermediate, were largely unaffected by exposure to higher temperature (Fig. 4B).

**Figure 4.**
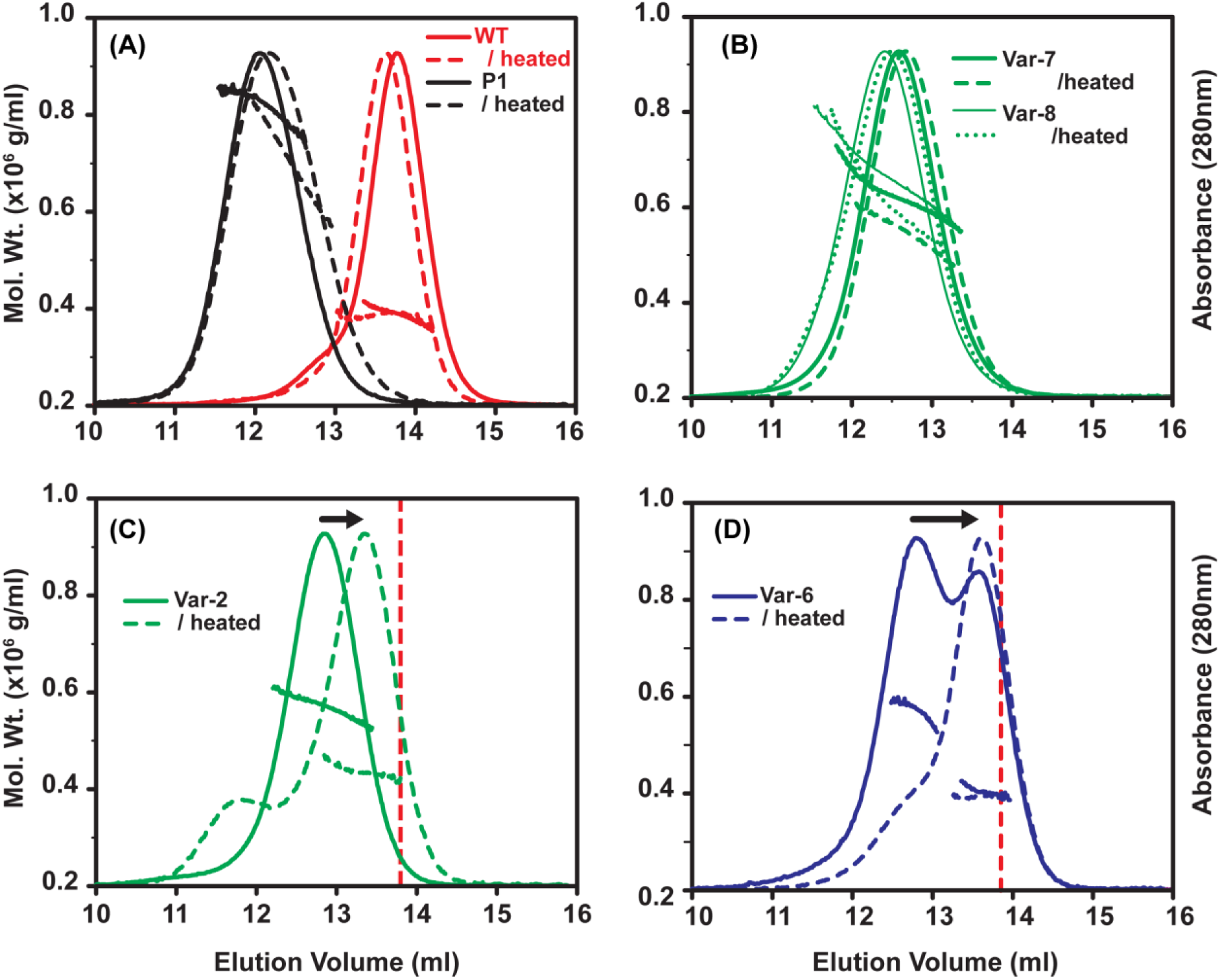
Temperature-induced oligomer rearrangement in Hsp16.5 variants. Variants were analyzed by SEC-MALS before (continuous line) and after incubation at 68 °C for 30 min (dashed line). A-B, the 24-subunit WT, the 48-subunit P1-insert, and the intermediate-size Var-7 and Var-8 oligomers were not susceptible to temperature-induced structural rearrangement. C, In contrast, Var-2 oligomers reassembled as a smaller oligomer upon heat treatment. D, incubation of Var-6 induced an increase of the WT-like 24-subunit arrangement and a reduction of the intermediate-size oligomer. The dotted vertical lines in C and D benchmark the WT 24-subunit elution peak.

In contrast, the intermediate oligomers formed by the poly-Ala Var-2 eluted from the SEC column with a larger retention volume (Fig. 4C). The temperature-induced change in average molar mass of Var-2 is consistent with a rearrangement into a smaller oligomer. Reduced oligomeric stability was exacerbated for the highly truncated P1 constructs of Var-5 and Var-6, which readily partition between two discrete populations. Incubation at high temperature resulted in depletion of the 36-subunit peak while simultaneously increasing the population of 24-subunit oligomers (Fig. 4D). Presumably the free energy of transition between the 36-subunit and 24-subunit oligomers at ambient temperature is sufficiently large to enable isolation of the two forms by SEC. Upon re-injection, the isolated 36-subunit and the 24-subunit oligomers of Var-6 eluted as distinct symmetrical peaks, did not scramble during extended storage at 4 °C (Fig. 5A and Fig. S7) and their molar masses agreed with their equilibrium populations (Fig. 5A). However, after incubation at 68 °C, the 36-subunit oligomeric pool of Var-6 shifted and converged upon the 24-subunit form according to SEC-MALS. This 24-subunit oligomer did not dissociate further with heat treatment, indicating that fundamental subunit interactions were preserved (Fig. 5B).

**Figure 5.**
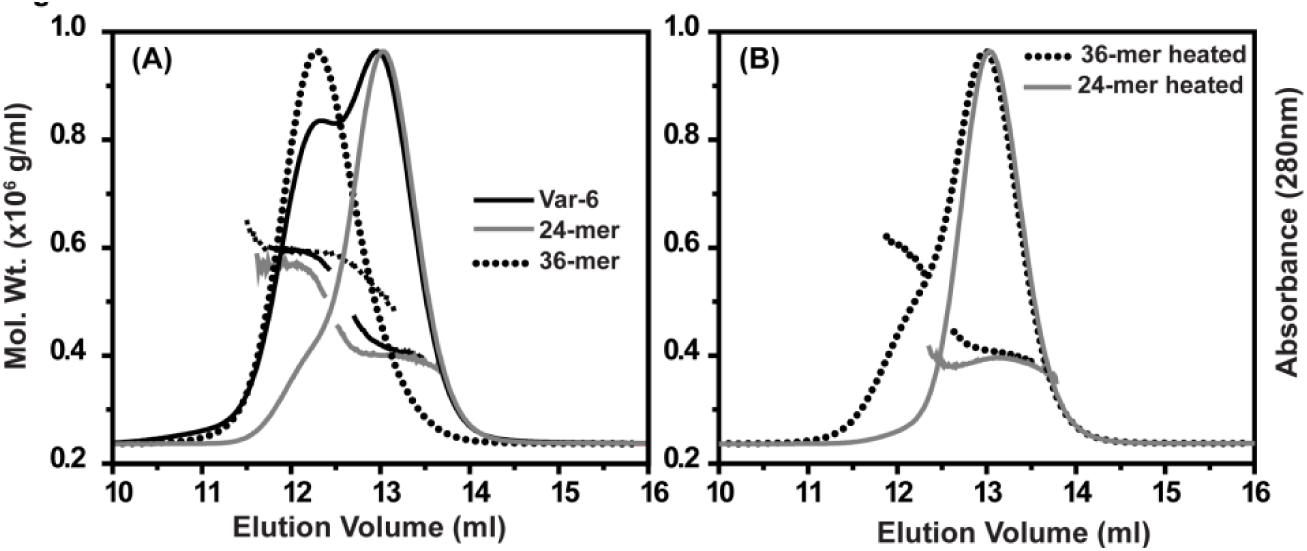
Isolation, characterization, and rearrangement of Var-6 oligomer subpopulations. A, 24- (gray trace) and 36-subunit (dotted trace) oligomers of Var-6 demonstrated similar MALS size distribution properties as the equilibrium populations observed prior to isolation by SEC. B, Incubation of the isolated 36-subunit oligomer (dotted trace) at 68 °C for 30 min induced collapse of the oligomer to a more thermostable 24-subunit arrangement (gray trace).

We further analyzed the oligomeric size distribution of four variants by BN-PAGE following heating at four different exposure times (Fig. 6). Var-5 and Var-6 showed a pattern consistent with that observed by SEC favoring a 24-subunit, WT-like oligomer. Remarkably, both Var-2 and Var-9 showed preferences for a species larger than the WT after heating, suggesting that full-length insertion is less compatible energetically with the 24-subunit assembly. BN-PAGE reveals the population of oligomers with an apparent composition of approximately 30 subunits. While the resolution of this technique does not allow for accurate determination of the number of subunits, the data shows that Hsp16.5 can occupy a ladder of different size oligomers consistent with the results from native state MS.

**Figure 6.**
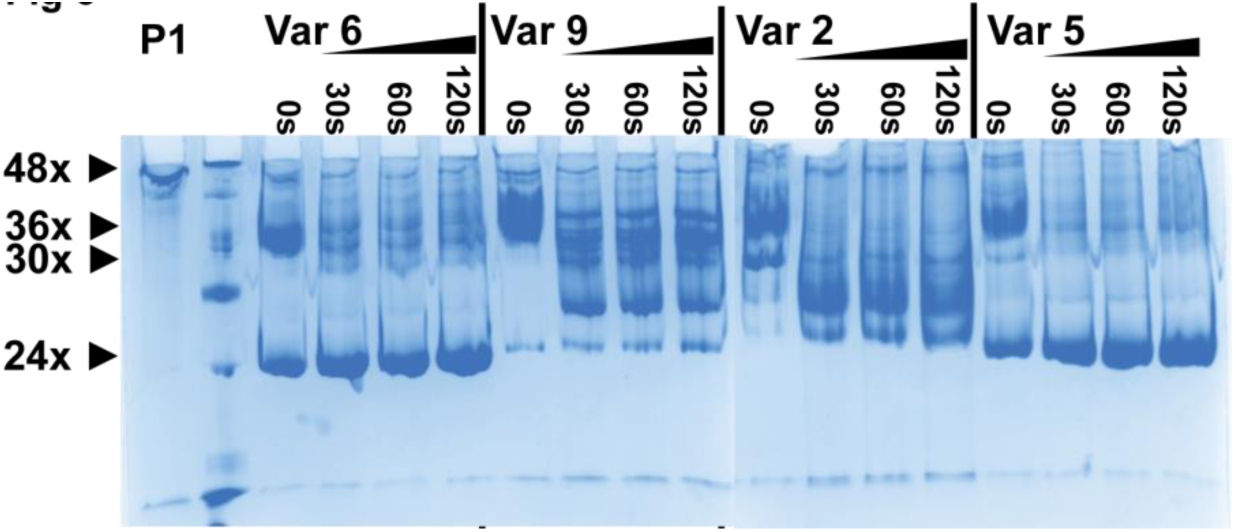
Temperature-induced subunit-rearrangement in Hsp16.5 variants. 30µg proteins were incubated at 68º C for 0, 30, 60, and 120 min, and were resolved on 7.5% polyacrylamide gels.

Together, these results suggest that although intermediate-sized assemblies are likely populated in the expansion into the ordered 48-subunit oligomer, these oligomers have lower stability than the 24 or 48 subunit oligomers. Interpreted in the context of the two-configuration model of the C-terminal tail, the striking differences in behavior between the variants upon heat treatment likely originate from the plastic interactions of the C-terminal tail with the α-crystallin domain of a neighboring subunit. Therefore, truncation of the P1 insertion, or altering its inherent flexibility (Var-2), compromises the adaptive response of the C-terminal tail and the ability to form stable, large oligomers.

### Heterogenous expansion enhances client-protein binding

A hallmark of the expanded Hsp16.5-P1 48-subunit oligomer is an increase in binding of client proteins relative to WT Hsp16.5, presumably due to greater accessibility of the high affinity binding sites within the N-terminal domain^47^. Likewise, a measurable increase in binding affinity would be anticipated for P1 variants that shift the oligomeric population toward larger structures relative to WT.

To characterize the binding efficiency of Hsp16.5 WT and P1 variants, we employed a well-established assay wherein a thermodynamically-destabilized model substrate (T4 lysozyme L99A, T4L-L99A) is titrated with increasing concentrations of the chaperone^53,54^. To monitor formation of a stable chaperone-substrate complex, T4L-L99A was labeled with the Cys-specific probe monobromobimane at residue T151C. Changes in bimane fluorescence anisotropy were observed upon addition of various molar ratios of Hsp16.5 variants, indicating the formation of a complex between Hsp16.5 and T4L (Fig. 7)^35^. Previous applications of this assay that utilize detection of fluorescence intensity revealed the two-mode nature of chaperone binding characterized by low and high affinity modes each manifested by distinct number of binding sites and fluorescence intensity values. Since probe anisotropy is similar in both low and high affinity binding modes, we performed the assay under conditions in which the high affinity mode dominates to reduce convolution with low affinity binding parameters^55^. The experimental binding curves were fit using a non-linear least squares approach assuming a single-mode binding model to obtain an apparent *K*_D_. The number of binding site was set to 0.5 T4L molecule per Hsp16.5 subunit to account for contribution by the low affinity mode for some of the variants.

**Figure 7.**
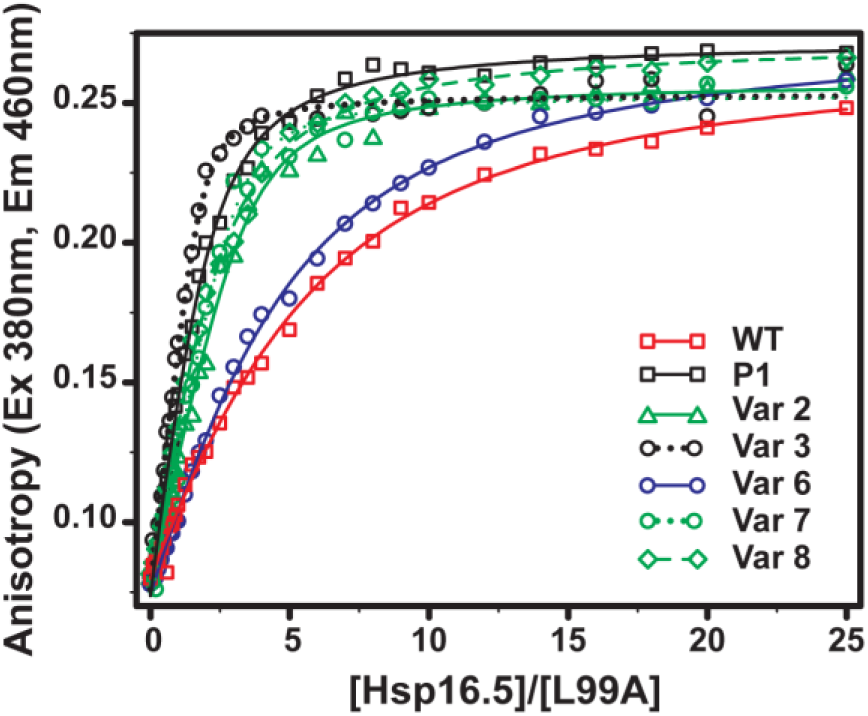
Binding isotherms of selected Hsp16.5 variants to bimane-labeled T4L-L99A. An increase in binding affinity correlates with variants that form large oligomers relative to WT. T4L (5μM) was incubated with increasing molar concentration of sHSP variants for 2hrs at 37 °C in pH 7.2 buffer. The solid lines are non-linear least-squares fit to a single-mode binding model, and the *K*_D_ for each variant is reported in Table 4.

Binding isotherms of WT Hsp16.5 and P1 variants to T4L-L99A are compared in Fig. 7 and the parameters are shown in Table 4. For all constructs, a monotonic increase in bimane anisotropy was observed followed by a region of saturation at high Hsp16.5 concentrations as expected. Consistent with previous studies, insertion of the full P1 peptide increases binding affinity by an order of magnitude^47^. Furthermore, in most cases, T4L-L99A was bound to a higher level by Hsp16.5 P1 mutants than by the WT revealing a correlation between apparent *K*_D_ and the propensity to form larger oligomers. For instance, Var-7 and Var-8, which form oligomeric ensembles clustered around 36-subunit sized assemblies, demonstrate a similar *K*_D_ to Hsp16.5-P1. In contrast, Var-6, which possesses a substantial fraction of 24-subunit oligomers, binds with an affinity similar to Hsp16.5 WT. Together with the structural analyses, these results reinforce the direct link between oligomer expansion and chaperone activity. The finding that Var-7 and Var-8 have similar binding characteristics to Hsp16.5-P1 suggests equivalence between polydispersity and expansion in increasing affinity to substrates.

**Table 4.** Dissociation constants of Hsp16.5 variants to T4L-L99A

| HSP variant | K<sub>D</sub> (μM) |
| --- | --- |
| WT | 11.96±0.84 |
| P1 | 1.28±0.09 |
| Var-2 | 2.65±0.29 |
| Var-3 | 0.19±0.04 |
| Var-6 | 9.47±0.48 |
| Var-7 | 1.58±0.23 |
| Var-8 | 2.22±0.13 |

5μM monobromobimane labeled T4L-L99A was titrated with increasing concentrations of sHsp at 37 °C in pH 7.2 buffer. The number of binding site binding site parameter, n, was constrained to 0.5 in all fits.

### Role of the PLPP motif in modulating the polydispersity of human Hsp27 oligomers

While the crystal structure of the expanded Hsp16.5-P1 oligomer did not resolve the P1 peptide, the N-terminus of the P1 peptide contains a low complexity sequence, PLPP, that occurs in proteins at higher frequency than expected and generally adopts a conserved rigid loop structural motif^56^. Given the results above suggesting a critical role of the PLPP motif in conferring the properties of the P1 peptide, we explored its importance in modulating the equilibrium dissociation of human Hsp27 oligomers. Hsp27 undergoes concentration-dependent dissociation from a polydisperse oligomer to a functionally-active dimer, and the two oligomeric states of Hsp27 exist in equilibrium at physiological pH^41,57,58^. This equilibrium of Hsp27* is illustrated by the series of analytical SEC of the purified protein at progressively lower concentrations (Fig. 8A). Injected at or above 500µg/ml concentration, Hsp27* eluted from the SEC column at retention volumes expected for the heterogeneous large oligomer. Progressive dilution of the protein led to the appearance of a peak corresponding to the dimer, which predominates at 10µg/ml of injected concentration. Stabilization of the dissociated dimer was observed at all concentrations for the Hsp27*-D3 mutant (S15D/S78D/S82D), which mimics the active, fully-phosphorylated state (Fig. S6)^59,60^. Deletion of the P1 peptide in both the Hsp27 and Hsp27-D3 has been shown to favor the larger oligomeric assembly, suggesting that the P1 peptide is important for the structural rearrangements mediating dynamic disassembly of larger oligomers^36^. Whether the diminished dissociation of the Hsp27 oligomer is due to the truncated P1 spacer or due to the unique flexible elements of the P1 sequence is not clear.

**Figure 8.**
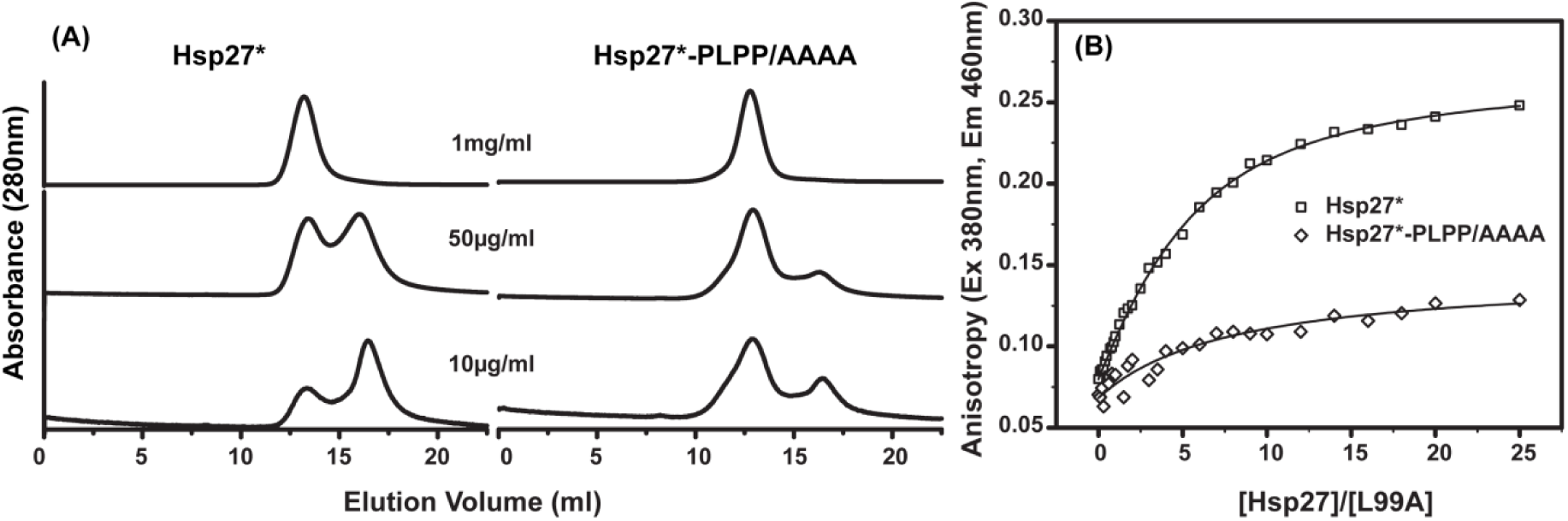
Impact of the PLPP motif on the assembly of Hsp27* oligomers and client protein binding. A, Concentration-dependent dissociation of the Hsp27* oligomer into the functionally-active dimer is disrupted when the PLPP sequence is substituted with a tetra-alanine peptide. B, Binding to bimane-labeled T4L-L99A is highly attenuated for Hsp27*-PLPP/AAAA mutant, which correlates with reduced propensity to form the dimeric species. T4L (5μM) was incubated with increasing molar concentration of Hsp27* for 2hrs at 37 °C in pH 7.2 buffer

To probe the importance of the PLPP structural unit in the dissociation equilibrium of Hsp27*, we substituted the sequence with tetra-alanine (AAAA). The mutant Hsp27*-PLPP/AAAA eluted from the SEC column at the retention volume identical to that of Hsp27* when injected at 1mg/ml and 500µg/ml concentrations (Fig. 8A). However at less than 100µg/ml injected concentrations, the peak corresponding to the dimer was less populated in the substitution mutant than in the WT. Thus replacement of PLPP mimics to a large extent the deletion of P1, suggesting a critical role in the modulation of oligomer dissociation. At the level of chaperone function, impaired transition to the dimeric species correlated with a loss of high affinity substrate binding in the PLPP/AAAA mutant (Fig. 8B).

### Concluding remarks

Sequence analysis of the P1 peptide reveals a remarkable divergence across species (Fig. 9). Indeed, only *H. sapiens*, *P. troglodyates* and *M. mulatta* have identical P1 sequences. Even in the evolutionary-close *Mus musculus*, P1 has undergone substantial sequence changes. Thus, our results demonstrating profound effects on polydispersity and chaperone-like activity as a consequence of P1 sequence changes predict a vastly different dynamic behavior of Hsp27 across the evolutionary spectrum.

**Figure 9.**
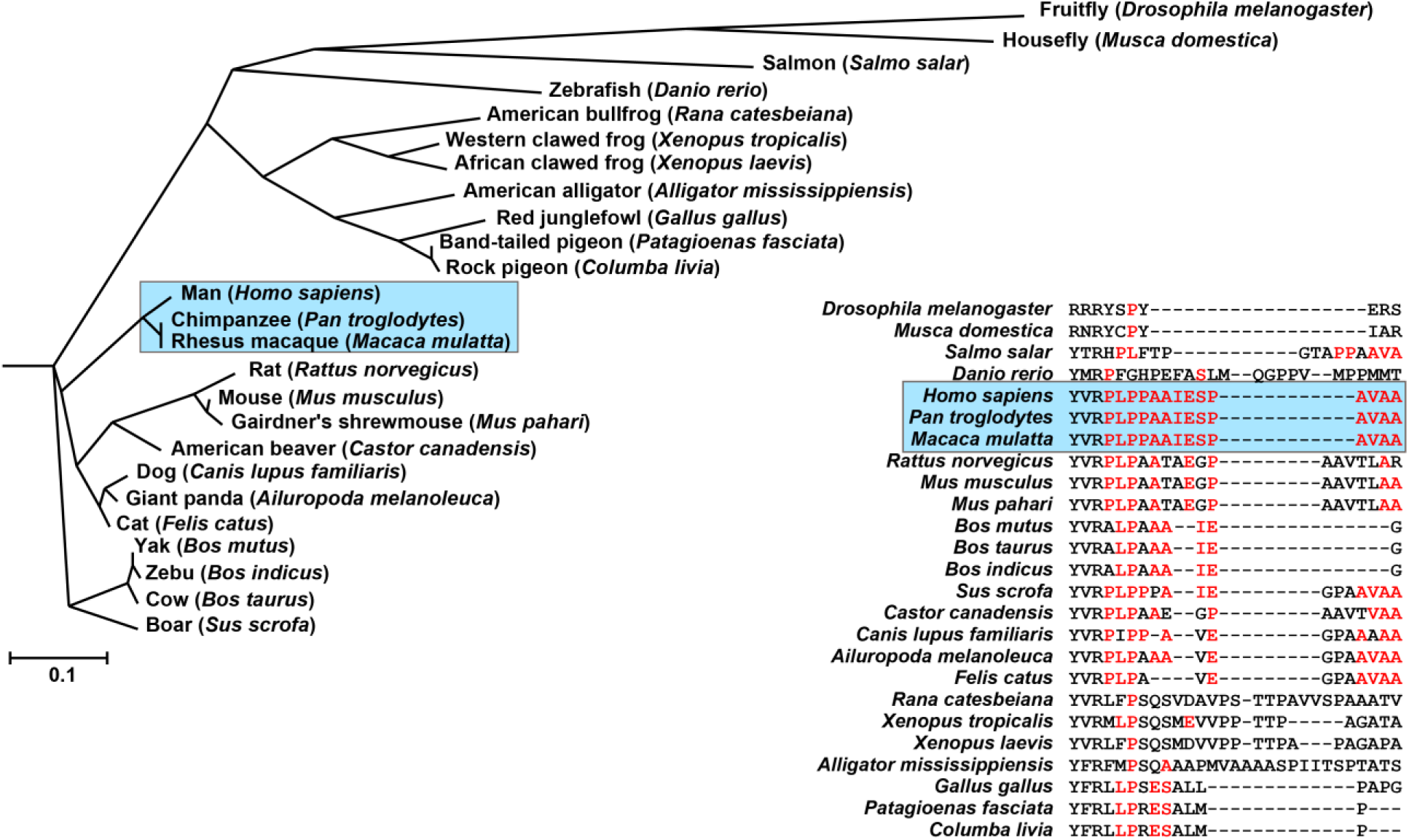
Sequence alignment of the P1 peptide and the evolution of Hsp27/HspB1. The P1 sequence was queried through the protein suite of the BLAST web engine. Among the Hsp27/HspB1 sequences, those representing diverse taxa were multisequence aligned by the Clustal Omega program. The phylogenetic tree was generated from the alignment by Neighbour-joining clustering and the cladogram was visualized using the Phylodendron web utility. Primates, highlighted by the box in the cladogram, have identical P1 sequence.

In a more general context, our results suggest structural and functional implications for the modular architecture of sHSP wherein the highly conserved modules, the α-crystallin domain and C-terminal tail are paired with a hydrophobic, sequence variable N-terminal domain. We propose that this evolutionary strategy enables diversity in the size and dynamics of sHSP oligomers as determined by unique sequences between the N-terminal and α-crystallin domains. The oligomer structure of *Triticum aestivum* Hsp16.9 uncovered the flexibility of the interactions of the C-terminal tail with the α-crystallin domain as one of the elements underlying alternative packing within the same oligomer^23^. Crystal and cryoEM structures of Hsp16.5-P1 demonstrated that the conformation of the C-terminal tail is structurally coupled to the sequence of the N-terminal domain^45^. The results presented here complete this model by establishing that polydispersity is intrinsically encoded in the architecture of the sHSP and may rationalize the extensive sequence divergence of the N-terminal domain.

While these design principles underpin conserved elements of the mechanism of sHSP chaperone activity, they also allow tuning and regulation of affinity and binding capacity to client proteins. Comparison of the structural and functional properties of archaeal Hsp16.5 and mammalian Hsp27 underscores conserved features of chaperone-like activity mediated by two distinct mechanisms of oligomer structure. Similar to the dynamic dissociation of large and polydisperse Hsp27 oligomers, the oligomer expansion model of activation is triggered by structural determinants encoded by sequences in the N-terminus and the consequent repacking of the C-terminal tail. On the other hand, the heterogeneity of client proteins and the need for tight regulation of chaperone activity necessitated the evolution of a complement of human sHSP tailored for the specific demands of cells and tissues. Similarly, physiological requirements likely led to the emergence of species-specific forms such as mouse αA^ins^-crystallin where a peptide insertion leads to an isoform of αA with high affinity to client proteins^35^. The interplay of these evolutionary forces accounts for the lack of strict conservation of sHSP oligomeric structure and the observed diversity in their size, symmetry and functional activity.

## Abbreviations

sHSP: Small heat-shock proteins
SEC: size exclusion chromatography
MALS: multi-angle light scattering
BN-PAGE: blue native gel electrophoresis
EM: electron microscopy
TEM: transmission electron microscopy
MS: Mass Spectrometry
T4L: T4 lysozyme
WT: Wild Type
MOPS: 4-morpholinepropanesulfonic acid
EDTA: Ethylenediaminetetraacetic acid
DTT: Dithiothreitol
Tris: 2-Amino-2-(hydroxymethyl)-1,3-propanediol
IPTG: isopropyl-β-D-thiogalactopyranoside
DEAE: diethylaminoethanol
IPTG: isopropyl-β-D-thiogalactopyranoside
SDS-PAGE: 
RI: refractive Index

## Material & Methods

### Cloning and mutagenesis

Site-directed mutagenesis of T4L-L99A and insertion of Hsp27 P1 peptide (the 14-amino acid peptide from residues 57-70 of human Hsp27) in the Hsp16.5 gene between residues 33 and 34 is described previously^47,48,61^. The variants of Hsp16.5-P1 with varying lengths of the insertion peptide based on the P1 sequence: Var-3 to Var-10 were generated by the QuikChange method (Stratagene) on the Hsp16.5-P1 template. The tridecamer alanine insert Var-2 was generated by overlap extension PCR as detailed elsewhere^62^. Hsp27 was cloned and the cysteine-less Hsp27, referred hereinafter as Hsp27*, was generated by the site directed mutagenesis of native cysteine at position 137 with alanine, as described previously^41^. It has previously been shown that the cysteine to alanine substitution does not affect the chaperone efficiency of Hsp27* in aggregation assays and equilibrium binding to T4L^22^. The Hsp27*-PLPP/AAAA mutant was generated by substituting proline – lysine – proline - proline residues at positions 57-60 to alanines by QuikChange. The phosphorylation mimicking triple mutant Hsp27*-D3 was generated by site-directed mutagenesis S15D, S78D, and S82D on the cysteine-free Hsp27* as described previously^57^. The plasmid constructs were confirmed by DNA sequencing.

### Expression, purification, and labelling of T4-Lysozyme

T4L-L99A mutant was expressed, purified, and labelled with monobromobimane as described previously^53^. Briefly, *Escherichia coli* cultures inoculated from overnight seeds were grown by shaking at 37°C until OD600 ~ 1 and induced by 0.4 mM isopropyl-β-D-thiogalactopyranoside (IPTG). After 3 hr induction at 30°C, the cells were harvested by centrifugation, resuspended into the lysis buffer (25mM MOPS, 25mM Tris, pH 7.6; 1mM EDTA, 0.02% (w/v) NaN_3_, 10 mM DTT), disrupted by sonication and centrifuged (15000 xg). T4L-L99 was purified by cation exchange on Resource S column by eluting with linear gradient of 1M NaCl. The eluted protein was incubated with 10-fold excess of monobromobimane for two hours at room temperature, followed by overnight incubation at 4°C. Unbound label was removed in the subsequent size exclusion chromatography (SEC) on a Superdex 75 10/300 GL column (GE Healthcare Life Sciences).

### Expression and purification of sHSPs

Hsp16.5 and the variants of Hsp16.5-P1 were expressed and purified by anion exchange, hydrophobic interaction, and size exclusion chromatography (SEC) as described previously^47,61,63^. Briefly, *Escherichia coli* cultures inoculated from overnight seeds were grown by shaking at 37°C until OD600 ~ 0.9 and induced by 0.4 mM IPTG. After 3 hr induction at 37°C, the cells were harvested by centrifugation (15000 xg), resuspended into the lysis buffer (20 mM Tris, pH 8.0; 100mM NaCl, 0.1 mM EDTA, 0.02% NaN_3_), disrupted by sonication, and the nucleic acids precipitated by polyethyleneimine (0.06% w/v). After clearing up the cell debris by centrifugation, Hsp16.5 variants were purified by anion exchange on a source Q column by eluting with linear gradient of 1M NaCl. Ammonium sulfate was added to a final concentration of 1 M, and the samples were loaded on a phenyl sepharose column and eluted with reverse gradient of 1 to 0 M ammonium sulfate. Proteins were finally purified by size exclusion (SEC) chromatography on a Superdex 200 10/300 GL or a Superose 6, 10/300 GL (GE Healthcare Life Sciences).

Hsp27* and Hsp27*-PLPP/AAAA were purified by sequential anion exchange and size exclusion chromatography as described previously^41^. Briefly, *Escherichia coli* cells were expressed by autoinduction for 12 h at 30 °C^64^. The cells were harvested by centrifugation and resuspended in lysis buffer (20 mM Tris, pH 8.0; 75 mM NaCl, 1 mM EDTA, and 10 mM DTT). The resuspension was sonicated and DNA was precipitated with polyethyleneimine (0.05% w/v). The lysate was cleared by centrifugation (15000 xg) and purified by anion exchange chromatography on Source Q column. Proteins were finally purified by size exclusion (SEC) chromatography on a Superose 6, 10/300 GL column (GE Healthcare Life Sciences).

*Escherichia coli* cells expressing Hsp27*-D3 by autoinduction for 12 h at 30 °C were lysed in the buffer containing 20 mM Tris, 1 mM EDTA, and 10 mM DTT and the supernatant as above was loaded on DEAE Sepharose column. Ammonium sulfate was added to the final concentration of 0.5 M, and protein was further purified from a phenyl sepharose column with reverse gradient of 0.5 to 0 M ammonium sulfate. Proteins were finally purified by size exclusion (SEC) chromatography on a Superose 6, 10/300 GL column (GE Healthcare Life Sciences).

All proteins were eluted from SEC columns in the SEC buffer containing 9mM MOPS, 6mM Tris, 50mM NaCl, 0.1mM EDTA, and 0.02% (w/v) NaN_3_ at pH 7.2. Protein concentrations were determined from absorbance at 280nm based on the extinction coefficients calculated by the ProtParam tool at the ExPASy webserver^65^. The homogeneity of all purified proteins was confirmed by SDS-PAGE, stained by Coomassie staining.

### Analytical Size Exclusion Chromatography

Analytical SEC on purified HSPs was performed on Superose 6, 10/300 GL column (GE Healthcare Life Sciences) equilibrated in the SEC buffer described above in an Agilent HP1100 HPLC system. Elution profiles of the proteins were monitored by absorption at 280nm wavelength using a UV detector and by the tryptophan fluorescence (Excitation: 295nm; Emission 330nm) using a fluorescence detector. Proteins were injected from 100µl loop at the indicated concentrations at an isocratic flow rate of 0.5ml/min.

### Molar Mass Determination

Molar mass of the proteins were determined by the multi-angle laser light-scattering detector (Wyatt Technologies) connected in-tandem to a refractive Index (RI) detector (Agilent). 100µl of each protein at 1mg/ml concentrations was injected by an Agilent HP1100 HPLC system on a Superose 6 column equilibrated in the SEC buffer at the isocratic flow rate of 0.5ml/min. The elution of the proteins was also monitored by an absorbance detector. The molar mass of the proteins were calculated by the Astra software (Wyatt) from the concentration of proteins derived from the RI signal based on constant dn/dc (0.185 ml/g).

### Blue-Native PAGE

The oligomers were resolved at pH 7 by electrophoresis on hand casted imidazole tricine polyacrylamide gels of the indicated acrylamide concentration in presence of 0.02% Coomasie Blue G-250 (Serva)^66^. Electrophoresis was performed in a cold room (BioRad MiniProtean 3 cell) at 100V until the samples entered the resolving gel, followed by constant 150V (current limited to 15mA) for ~2 hours. The gels were visualized by standard Coomasie staining.

### Subunit rearrangement

Proteins diluted to 1mg/ml in the SEC buffer were incubated in Eppendorf tubes at 68 °C for 30min, unless specified otherwise, in a water bath to stimulate subunit rearrangement. The thermally induced subunit rearrangement was quenched by transferring sample tubes to ice-water slush.

### Binding of T4L-L99A to Hsp16.5 and Hsp27* variants

Solutions of monobromobimane-labelled T4L-L99A at final concentration of 5µM with increasing molar excess of Hsp16.5 or Hsp27* variants diluted in the SEC buffer were incubated at 37 °C for 2 h. The fluorescence anisotropy was measured in a SynergyH4 microplate reader (BioTek) at 37 °C. The monobromobimane label was excited by 380nm filter (bandpass 20nm) and the emission fluorescence in the horizontal and vertical polarizations was read by 460nm filter (bandpass 40nm). The binding isotherms were analyzed on Origin 8 (OriginLab Corporation, Northampton, MA) by the nonlinear least-squares fits using the Levenberg–Marquart method.

### Electron Microscopy

10 nM droplets of proteins were applied for two minute to freshly glow discharged solid carbon support on copper 400 mesh TEM grids. Excess water was blotted away and replaced with pH neutralized 0.75% uranyl formate solution, then allowed to dry. Prepared grids were visualized at a nominal magnification of 67,000X on an FEI Tecnai T12 operating at 120 KeV. Digital images were captured using the Semi-Automated Microscopy package^67^ on a Gatan US1000 CCD camera. Images were normalized and individual HSP particles were selected from micrographs of each mutant variant using the EMAN Boxer routine^68^. The total number of particles for each sample is as follows: P1 = 6853, Var2 = 2550, Var3 = 2747, Var4 = 4987, Var5 = 4756, Var6 = 2846, Var7 = 5770, Var8 = 22040, Var9 = 9834 particles. Previously determined Hsp16.5 24mer, 36mer and 48mer complexes were used as starting models for each mutant variant in multirefine.py subroutine from EMAN. Each particle set was individually refined against the three starting structures to determine the relative distribution of assembly states for each mutant. Post-multirefine.py bins for each of the three resulting structures were reclassified in EMAN refine2d.py to confirm the sorting process.

### Native mass spectrometry

Native mass spectrometry was performed as described elsewhere for oligomeric identification and quantification^69^. Briefly, ions were generated from a 200 mM NH4OAc solution and introduced into a Q Exactive Plus hybrid quadrupole-Orbitrap mass spectrometer (Thermo Fisher Scientific, Bremen Germany) modified for the transmission, selection and detection of high mass ions^70^. All spectra were acquired in “EMR Mode” and a pulser mode of 50. Ions were generated in positive polarity from a nanospray source using gold-coated capillaries prepared in-house. Transfer capillary temperature was (100-120 °C). The injection flatapole was modified for in-source trapping and a desolvation voltage of -100 V applied. No in-source activation was used. Ions were trapped in the HCD cell before being transferred into the C-trap and Orbitrap mass analyser for detection. Transient times were 64 ms and AGC target was 1×10^6^. The collision gas was nitrogen and UHV pressure was maintained at approximately 1×10^−9^ mbar. Spectra were acquired with 10 microscans, averaged over a minimum of 100 scans and with a noise level parameter set to 3. For MS/MS experiments the HCD activation voltage was increased to between 150-250 V to achieve full or near-complete dissociation of the selected precursor. Tandem MS was performed across the whole oligomeric range where sufficient ion intensity allowed.

Intact oligomeric distributions were fitted to spectra, where possible, using mass spectrometry deconvolution software, Unidec, as described previously^71^. All spectra underwent minimal curved background subtraction with a value of 100, except variant 4 which is shown with a value of 20^72^. CID spectra were assigned using Unidec software and the precursor identity determined manually. Estimates for the relative abundance of each oligomeric cluster (24, 28-38, 48) were achieved through integration of the intact MS spectrum over the m/z regions shown in Fig.S3. Relative abundances for each individual oligomeric state, including discrimination by truncation number, were determined for variants 2, 4 and 8 by direct fitting of oligomeric species to the intact spectrum using Unidec. Resulting fit distributions were consistent with CID spectra for the corresponding range. The relative abundance for the 28-38 subunit region for variants 5 and 6 was calculated from the reconstruction of tandem MS species weighted by the integral of the spectrum across the quadrupole selection window. This was then scaled relative to the well resolved 24 subunit region in each case. The 36 subunit cluster was then generated for comparison through summation of the oligomeric states 28-38. All deconvoluted data was scaled according to the reciprocal of the charge state to account for the magnitude of charge on the detector signal.

## Acknowledgment

This work was supported by National Institutes of Health Grant R01-EY12018 to H.S.M. S.A.C. and J.L.P.B. thank the Biotechnology and Biological Sciences Research Council (BB/L017067/1) and Waters Corp for an iCASE studentship. We thank Joseph Gault (Oxford) for useful discussions and advice

## Author Contributions

Conceptualization, S.M. and H.S.M; Validation, S.M., S.A.C., D.W., J.L.P.B., H.S.M.; Investigation, S.M, S.A.C, D.W., H.A.K.; Writing – Original Draft, S.M., S.A.C., D.P.C., H.S.M.; Writing – Review & Editing, S.M., S.A.C., D.W., D.P.C., P.L.S., J.L.P.B., H.S.M.; Visualization, S.M., S.A.C., D.W.; Supervision, P.L.S., J.L.P.B., H.S.M.; Funding Acquisition, S.A.C., J.L.P.B., H.S.M.

